# ScAPE: A lightweight multitask learning baseline method to predict transcriptomic responses to perturbations

**DOI:** 10.1101/2025.09.08.674873

**Authors:** Sergio Romero-Tapiador, Pablo Rodriguez-Mier, Martin Garrido-Rodriguez, Ruben Tolosana, Aythami Morales, Julio Saez-Rodriguez

## Abstract

Predicting biological responses to perturbations is essential for advancing drug development efforts. While high-throughput technologies have greatly enhanced our ability to study perturbation effects at the molecular level, accurately modeling these responses remains challenging due to the complex underlying molecular interactions, the vast dimensionality of cellular response, the limited amount of data, and technical limitations in data collection. Machine learning (ML) approaches have emerged as powerful tools for addressing these challenges. However, the observation in recent benchmarks that complex models do not consistently outperform simpler ones highlights the need for robust and efficient baselines for evaluating new methods. We present ScAPE (Single Cell Analysis of Perturbational Effects), an ML method for predicting drug-induced gene expression changes. ScAPE leverages only aggregated gene-level statistics across cell types and perturbations as input features, yet achieves good performance despite this simplicity. Its effectiveness was demonstrated in the Single-cell Perturbation Prediction NeurIPS 2023 challenge, where it ranked among the top-performing methods. We describe the core model and introduce extensions that enhance its utility as a baseline, including a multitask learning extension for simultaneous prediction of multiple transcriptional response metrics. A software API is provided to support adoption and integration into research pipelines. Benchmarking against other leading challenge entries and general-purpose ML approaches, using the original challenge metric, supports ScAPE as a simple, lightweight baseline for evaluating methods that predict perturbation effects at the gene-expression level. ScAPE is built using Keras 3, enabling support for different backends such as TensorFlow, JAX, or PyTorch, and freely available as an open-source package at github.com/scapeML/scape.

## 1 Introduction

Predicting how cells respond to chemical perturbations is a fundamental challenge in computational biology with important implications for drug discovery and personalized medicine. The ability to accurately infer cellular responses to compounds could reduce the time and cost of drug development by enabling *in silico* screening of promising candidates before costly experimental validation^1^. However, this task is exceptionally difficult due to the vast chemical space^2^ of potential perturbations, the complexity of cellular responses, and the limited availability of high-quality training data.

On the experimental side, single-cell RNA sequencing (scRNA-seq) has become a widely used approach for studying cellular responses to perturbations. Unlike bulk RNA sequencing, which averages signals across thousands of cells and obscures cell-to-cell variability, scRNA-seq captures transcriptional profiles at single-cell resolution, revealing heterogeneity that bulk measurements cannot detect^3,4^. This capability allows to profile millions of cells, making it possible to observe subtle responses to chemical treatments and other perturbations. The resulting datasets offer unprecedented opportunities for building predictive models of cellular behavior. Yet, predicting perturbation responses at the gene-expression level remains challenging. The perturbation space is vast—encompassing chemical compounds, genetic interventions (e.g., CRISPR, siRNA), ligands, biologics, and their combinations—each influencing gene expression in distinct ways and varying across biological contexts such as tissues, cell types, and individuals^5^. As a result, current predictive methods often fail to generalize beyond specific experiments, achieving only modest accuracy^6,7^.

In recent years, machine learning (ML)-based approaches have shown substantial promise in predicting gene expression responses, particularly in the context of perturbation modeling. Models such as scGen^8^, Dr.VAE^9^, or Compositional Perturbation Autoencoder (CPA)^10^ propose deep autoencoders to predict gene expression profiles under various perturbation conditions.

Additionally, recent advancements such as chemCPA^11^, which incorporates molecular structure information, and GEARS^12^, which integrates gene network structure into the model, illustrate different ways in which prior knowledge can be injected into the models as an attempt to improve generalization to unseen perturbations. Approaches grounded in representation learning and causal inference have also emerged as powerful alternatives to traditional methods, promising more accurate modeling of perturbation effects by disentangling latent biological factors and identifying causal relationships among variables^13–15^.

Despite these advancements, current models often fall short in terms of real-world performance: they struggle to generalize across diverse settings and frequently fail to outperform simpler baseline approaches, raising questions about whether their added complexity is justified^6,7,16,17^. To help address these challenges and push the field toward more robust and generalizable solutions, the Neural Information Processing Systems (NeurIPS) 2023 perturbation prediction competition provided a large-scale, biologically diverse dataset and a competitive benchmarking framework to assess model generalizability across compounds, cell types, and donors. The competition focused on predicting single-cell gene expression responses of human peripheral blood mononuclear cells (PBMCs) to small-molecule perturbations, and was hosted on Kaggle^1^ as part of the NeurIPS 2023 competition track^5^. The resulting dataset, termed Open Problems Perturbation Prediction (OP3), included scRNA-seq profiles of PBMCs treated with 144 compounds from the Library of Integrated Network-based Cellular Signatures (LINCS) Connectivity Map. The primary objective of the competition was to predict changes in gene expression, using signed − log_10_ multiple-testing-corrected p-values for each gene, cell type and drug, derived from differential expression analysis.

Here, we introduce the Single Cell Analysis of Perturbational Effects (ScAPE)^18^, an extension of one of our best-performing methods in the competition. ScAPE is a lightweight 18.9M-parameter neural network designed to predict gene expression changes for unseen drug–cell combinations. The model is not designed to extrapolate to entirely novel drugs or cell types; instead, it learns to generalize to new pairings where each component (drug or cell) has been observed in other contexts. Our method, which was awarded a prize in the competition for its strong predictive performance and well-motivated design, is presented here along with an extended analysis and enhancements. Conceptually, our approach draws inspiration from collaborative filtering in recommendation systems, where user preferences are predicted based on the preferences of similar users and the characteristics of similar items. In the perturbation prediction context, we hypothesize that a cell’s response to a drug can be predicted from two key pieces of information: (1) how that drug typically affects gene expression across different cell types (analogous to item characteristics), and (2) how that cell type typically responds to different perturbations (analogous to user preferences). These median-based signatures capture the essential biological signal while remaining robust to noise and batch effects. The simplicity of ScAPE makes it easy to adapt, modify, and train, offering a robust and accessible baseline for perturbation prediction tasks.

We also introduce an improved variant that incorporates multi-task learning and ensembling to jointly predict both p-values and *log*_2_ fold-changes (LFC), while significantly improving computational efficiency. Predicting the LFC provides the effect size of gene expression changes, while the p-value reflects their statistical significance, which can help the model distinguish robust signal from noise rather than focusing solely on large but potentially spurious effects.

We provide a comprehensive evaluation of both versions of the method, comparing them against other top-performing approaches from the NeurIPS 2023 competition, as well as TabPFN—a general-purpose foundation model for tabular data not specifically designed for perturbation prediction^19^. ScAPE is built using Keras 3, enabling support for different backends such as TensorFlow, JAX, or PyTorch, and freely available as an open-source package at github.com/scapeML/scape.

## 2 Material and Methods

### 2.1 Open Problems Perturbation Prediction (OP3) Dataset

The OP3 dataset was derived from experiments conducted on PBMCs from three healthy human donors, with gene expression measured 24 hours post-treatment. The experimental setup employed 96-well plates, incorporating two positive controls (dabrafenib and belinostat) and a negative control (DMSO), alongside wells dedicated to individual compound treatments. Each well contained PBMCs, which include various immune cell types such as T cells, B cells, and Natural Killer (NK) cells.

The dataset comprises differential expression (DE) profiles for 18,211 genes across multiple cell types in response to various compound perturbations, with DE quantified using signed log_10_ p-values (SLPs). The dataset was partitioned into training, public test, and private test sets, with only the training set accessible to participants.

The training set includes almost all compounds for T cells (CD4^+^, CD8^+^, and T regulatory cells) and NK cells. Additionally, a smaller subset, along with positive and negative controls, was provided for B and myeloid cells. For the test sets, the public test set contains data for 50 randomly selected compounds in B and myeloid cells, while the private test set comprises the remaining 79 compounds for these cell types.

In total, the OP3 dataset encompasses 614 unique cell type-compound combinations, posing significant challenges due to the high-dimensional feature space (18,211 genes) relative to the low-dimensional observation space. This complexity is further exacerbated by a low signal-to-noise ratio (SNR): although 7,699 genes (∼38%) explain 80% of the total SLP variance, the remaining ∼62% of genes display minimal variation.

### 2.2 Analysis of the OP3 Dataset

From more than ten million gene expression values, there are notable differences in the number of compounds available for each cell type. Specifically, there are 144 compounds for NK cells, CD4^+^ T cells, and T regulatory cells, 140 compounds for CD8^+^ T cells, and only 15 compounds for both B cells and myeloid cells. Most of the values cluster around zero, indicating that the majority of perturbations do not induce statistically significant changes in gene expression across most gene (**Figure 2a**).

**Figure 1:**
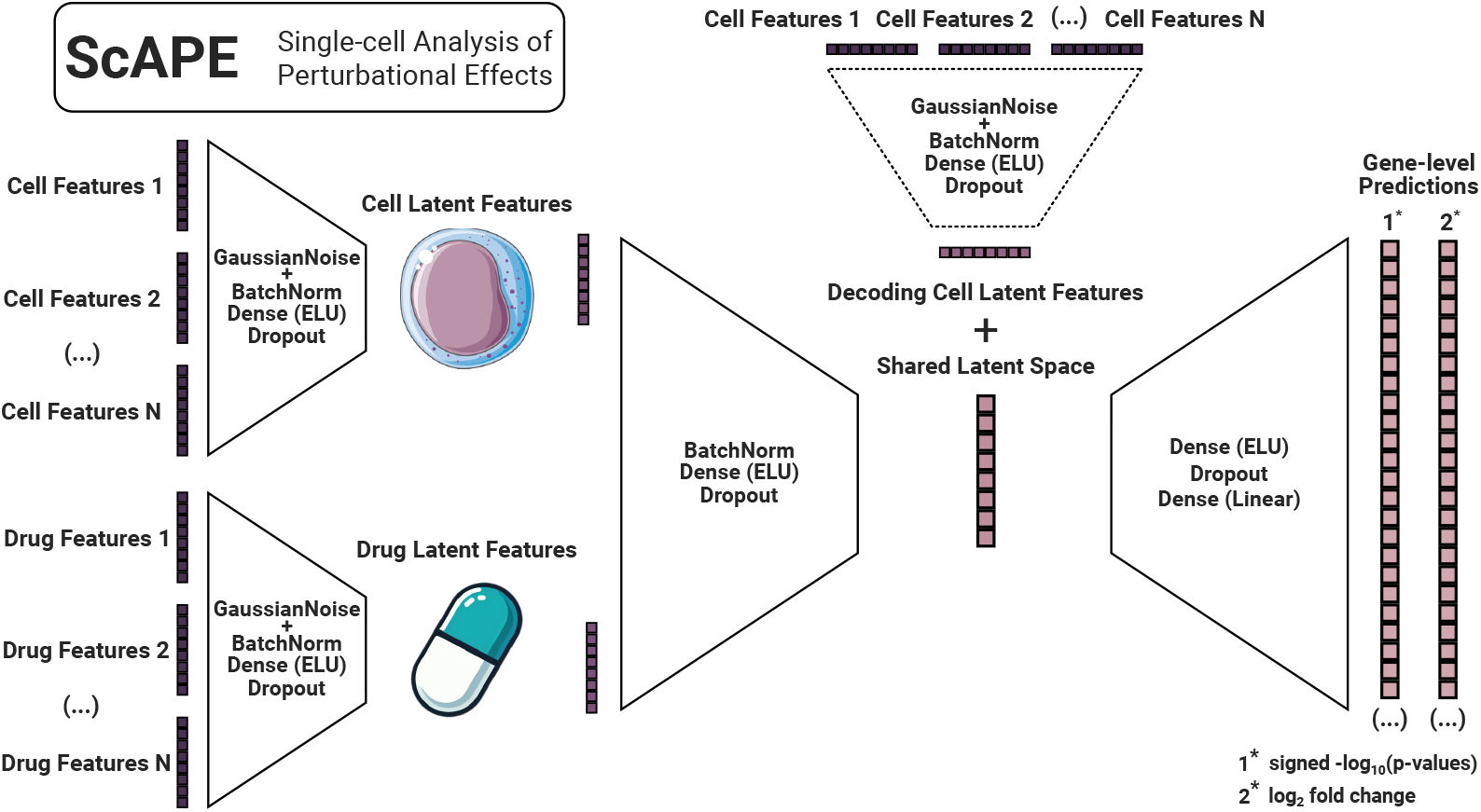
ScAPE (Single Cell Analysis of Perturbational Effects) architecture for perturbation prediction. Cell and drug features undergo Gaussian noise, batch normalization, and processing through dense exponential linear unit layers with dropout to produce latent representations. These are integrated into a shared latent space via an encoder, followed by a decoder that reintroduces cell latent features and generates gene-level predictions. The model outputs signed −*log*_10_(p-values) (SLPs), for the single-task, or both SLP and *log*_2_ fold changes for the multi-task approach.

**Figure 2:**
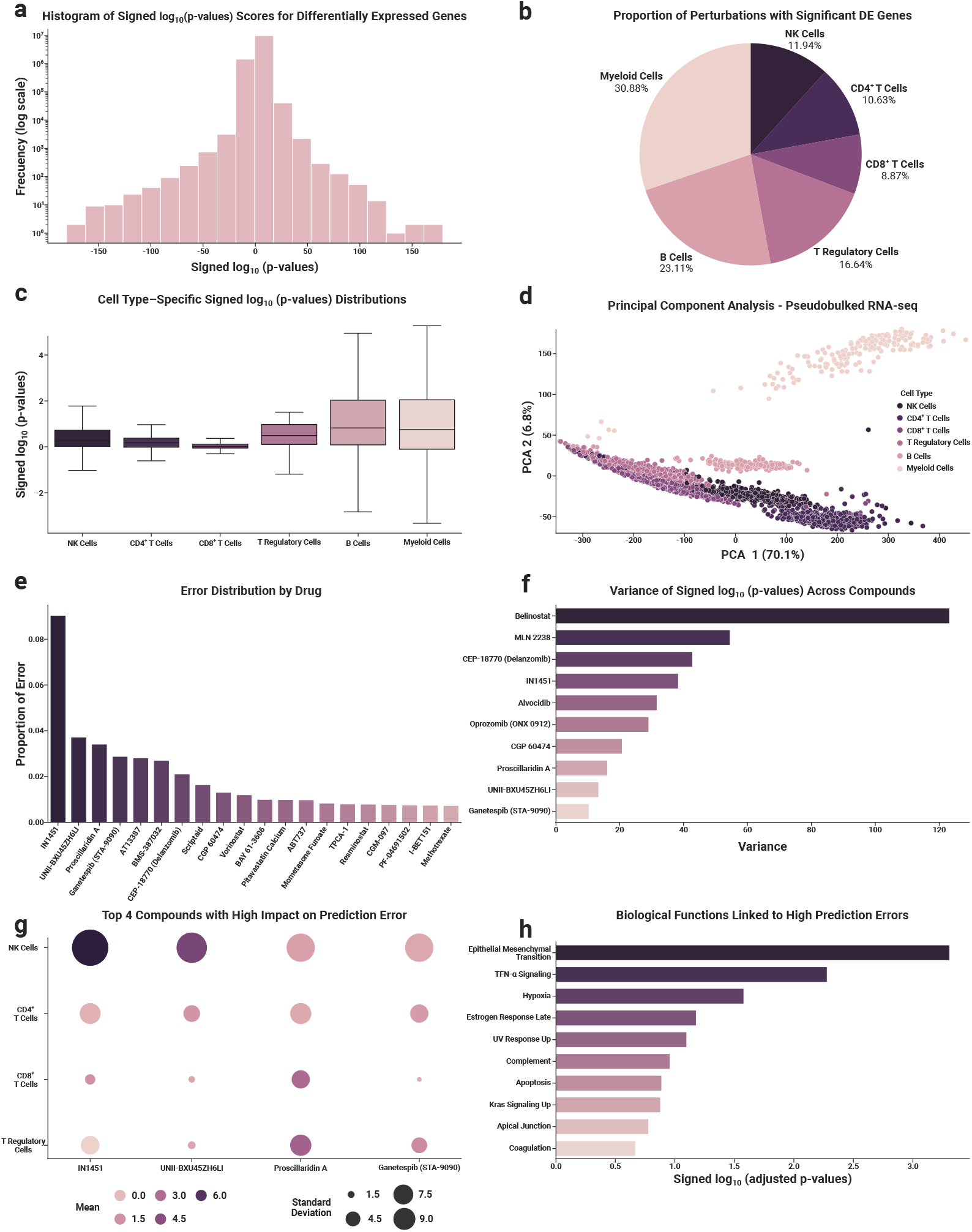
Overview of cell type, compound, and gene expression data along with corresponding analyses. a) Histogram of SLPs of the differentially expressed genes in the perturbation prediction dataset. b) Proportion of perturbations with at least one statistically significant deregulated gene, shown across cell types. c) Mean SLP values across different cell types. d) Principal component analysis (PCA) of pseudobulk RNA-seq data. e) Distribution of errors across drugs. f) Variance of SLP values across compounds. g) Top 4 compounds with the highest prediction errors across cell types. h) Enrichment analysis of genes with high prediction errors linked to key biological functions.

To better understand how transcriptional responses vary by cell type, we calculated the proportion of genes exhibiting statistically significant DE, defined as those with *SLP >* 1.3 (or equivalently *p <* 0.05). The results revealed substantial variability across cell types. Myeloid and B cells exhibited the strongest responses, with 30.88% and 23.11% of genes classified as differentially expressed, respectively. However, these two cell types were also exposed to the fewest compounds, suggesting that their elevated SLP proportions should be interpreted with caution. Among the cell types with broader experimental coverage, T regulatory cells showed the highest proportion of DE genes (16.64%), followed by NK cells (11.94%), CD4^+^ T cells (10.63%), and CD8^+^ T cells (8.87%). These findings align with the broader distribution patterns observed across the dataset (**Figure 2b**).

To complement this analysis, we assessed the overall distribution of SLP values to better characterize the intensity and consistency of transcriptional responses within each cell type. While most distributions were centered near zero, notable differences in spread and skewness emerged. CD8^+^ T cells displayed the narrowest distribution, suggesting consistently weak responses to perturbations. Conversely, B cells and myeloid cells exhibited wider spreads and higher median SLPs, indicative of stronger transcriptional shifts—though, as noted earlier, these results may be influenced by their limited compound exposure. The remaining cell types, including NK cells, CD4^+^ T cells, and T regulatory cells, showed moderate variability, reflecting intermediate response profiles that align with earlier observations on differential expression proportions (**Figure 2c**).

To visualize the structure of the dataset, we applied principal component analysis (PCA) to the pseudobulk RNA-seq data^20^. The resulting projection revealed that a small number of principal components account for the majority of the variance, with the first two alone explaining more than 75%. The projected data exposed clear differences in transcriptional profiles across cell types: myeloid cells exhibited the most distinct separation from the others, reflecting their unique expression landscape. B cells also displayed some divergence, though they remained more closely aligned with the broader distribution. The rest of the cell types—T cells and NK cells—were more tightly grouped, suggesting shared transcriptional features and more subtle inter-type variability (**Figure 2d**).

Among the tested compounds in PBMCs, we observed substantial differences in their effects on gene expression: epigenetic and proteasomal inhibitors such as belinostat (an HDAC inhibitor), MLN-2238 (ixazomib) and CEP-18770 (delanzomib) elicited highly variable transcriptional responses, reflecting broad chromatin de-repression and proteotoxic stress that activate hundreds to thousands of downstream genes in this immune cell context, whereas most other molecules, such as the 5-HT_1_A partial agonist buspirone, the tricyclic antidepressant protriptyline and the BMP-receptor kinase inhibitor K-02288, induced very few differentially expressed genes, consistent with their limited activity in PBMCs and their more focused receptor- or kinase-specific modes of action (**Figure 2f**).

To establish a predictive baseline, we first implemented a simple feed-forward neural network. Using this model, we conducted an initial exploratory experiment to characterize overall dataset variability and pinpoint key prediction challenges. We then examined the per-drug error distributions, which were largely uniform except for the first four compounds, which together accounted for 15% of the total error (**Figure 2e**). As anticipated, drugs that proved hardest to predict showed pronounced error shifts between training and test cell types. For example, IN1451, the single greatest source of error, elicited a strong response in NK cells but had minimal effects on CD4^+^ T cells, CD8^+^ T cells, and T regulatory cells (**Figure 2g**).

We further explored whether specific biological functions were more difficult to predict. An enrichment analysis was conducted on the top 5% of genes with the highest average error in our local setup, using MSigDB hallmarks^21^ and decoupleR^22^. This revealed that certain hallmarks, such as epithelial-mesenchymal transition and TNF-*α* signaling, contained a notable number of highly variable genes. This enrichment reflects underlying sensitivity differences. In drug-sensitive cells these pathways are strongly activated, driving inflammatory, stress or apoptotic responses, while in non-sensitive cells they remain near baseline. Such large expression differences across cell types result in higher prediction errors. By contrast, genes outside these sensitivity-linked programs exhibit smaller deviations from baseline and are therefore easier for the model to predict (**Figure 2h**).

### 2.3 Experimental Protocol

To establish a robust and consistent framework for evaluating and comparing ML models, we implemented a nested cross-validation strategy using clearly defined data partitions within the OP3 dataset. This structured approach enables fair model comparison and effective assessment of cell-type-specific predictions and model stability.

We partitioned the dataset initially into two main subsets: a training-validation dataset (∼ 80%) and a hold-out test dataset (∼ 20%). The training-validation dataset is subsequently split into distinct training and validation subsets, while the test dataset remains strictly isolated for final model evaluation. The outer validation loop, termed “cell-type cross-validation”, evaluates model performance across specific cell types. We used a 4-fold cross-validation scheme, where each fold designates one cell type (e.g., NK, CD4^+^ T cells, CD8^+^ T cells, or T regulatory cells) as the test set. The remaining cell types, along with a limited subset of the selected test cell type, form the training-validation dataset. These four cell types were selected based on their high representation within the dataset. In each fold, the test set includes 90% of compounds from the designated cell type.

Correspondingly, the training-validation set includes all cell type-compound pairs from other cell types not designated as the test cell type in the current fold. Additionally, the training-validation dataset includes 10% of compounds from the designated test cell type, ensuring representation of all cell types during training and all control compounds.

Within each outer fold, we apply a 5-fold inner cross-validation exclusively on the training-validation dataset to optimize model parameters. Here, the training data comprises all samples from B and myeloid cells, all samples from 20% of the compounds, and all but one sample from each of the remaining 80% of compounds. The validation data consists of the remaining single samples from each of the 80% of compounds not fully included in the training set. These validation samples are randomly selected from the three most numerous cell types in the current training-validation set.

This nested cross-validation design effectively addresses challenges associated with high dimensionality and low signal-to-noise ratios in biological data while preserving biologically meaningful structures. Combining an outer loop for evaluating cell-type generalizability with an inner loop focused on compound-based model tuning ensures comprehensive evaluation across diverse biological contexts. The allocation of a small proportion (10%) of compounds from the test cell type into the training-validation set guarantees cell type representation during training, mimicking the original setup of the competition, where we only observe a small number of perturbations for the cell types of interest. Final predictions for each cell type were generated through an ensemble method, calculating the median prediction across the five inner-loop folds.

### 2.4 ScAPE Model

**Figure 1** illustrates the ScAPE architecture, specifically designed for perturbation prediction. The final design consists of a simplified autoencoder with two variants, distinguished by the number of tasks they perform: single-task ScAPE and multi-task ScAPE. First, the single-task ScAPE produces a single output, predicting DE outcomes for cell type/drug pairs as SLP. Second, the multi-task ScAPE generates two outputs, i) the SLP outcomes (as in single-task ScAPE) and ii) LFC values, providing information about effect size. Both variants share the same core architecture, differing only in their output layers.

The strength of ScAPE lies in its use of two-dimensional median summaries for each gene, computed both across cell types and across drugs, as the sole input features. For every drug, we take the median SLP and the median LFC (in the multi-task mode) over all cell-type pseudobulk profiles; likewise, for every cell type we take those same medians over all drug pseudobulk profiles. These drug-level and cell-level effect vectors capture, respectively, the overall potency and statistical confidence of each compound at the gene level and the inherent context-specificity of the drug activity for each cell type.

To improve robustness, zero-mean Gaussian noise is added independently to each element of these vectors, which are then standardized via batch normalization. Each noisy, normalized vector is projected through a dense layer with exponential linear unit activation and dropout to produce latent drug and cell embeddings. Those embeddings are concatenated to form a shared latent representation that captures the joint drug–cell interaction space. In the decoding phase, a different cell-embedding vector is concatenated with the shared latent vector and passed through another ELU-activated dense layer with dropout, followed by a final linear layer that generates gene-level predictions.

In initial experiments, we explored a conditional Variational AutoEncoder (VAE) via a *β*-VAE formulation with an annealed Kullback–Leibler divergence, but found no improvement in predictive accuracy or generalization despite added complexity, similar to^10^. We therefore settled on a simpler, non-probabilistic neural network architecture for ScAPE.

Section 3.1 presents a detailed analysis of parameter tuning, including variations in the number of input genes, ensembling strategies, and loss function weight adjustments. The ScAPE architecture is implemented in the open-source ScAPE library^2^.

### 2.5 Benchmark

We benchmarked ScAPE against the two top-ranked submissions, as reported by “Open Problems in Single Cell Analysis” post-competition analysis (excluding ScAPE itself). We also included in the benchmark the recent TabPFN method, a powerful transformer-based foundation model, exemplifying a state-of-the-art, general-purpose predictive method not specifically developed for this competition.

- **NN retraining with pseudolabels**^3^: This method follows a two-stage approach. In the first stage, pseudolabels are generated using an NN optimized with Optuna, which produces predictions through a 7-model ensemble with custom cross-validation. In the second stage, the pseudolabels are incorporated into the training data, and predictions are refined using a 20-model ensemble, followed by median prediction clipping to ensure consistency. This system leverages pseudolabeling and ensembling to enhance prediction robustness.
- **Py-boost**^4^: The Py-boost approach utilizes four models. First, it combines gradient-boosted decision trees using t-statistics and PCA for dimensionality reduction. Additionally, ridge regression and k-nearest neighbors recommender systems are applied, framing the problem as collaborative filtering between cell types, compounds, and gene expression. The final model, ExtraTrees, employs fully grown decision trees with noise-enhanced target encoding for improved generalization. This ensemble approach effectively addresses the challenges of high dimensionality and low sample sizes.
- **TabPFN**^**23**^: The Tabular Prior-data Fitted Network (TabPFN) is a transformer-based foundation model pretrained on millions of synthetic tabular datasets. At inference time, it ingests a small labeled training set and unlabeled query points, and directly outputs approximate posterior predictive distributions. This enables rapid, few-shot classification and regression with strong performance on high-dimensional, low-sample-size tasks. By virtue of its learned inductive biases and ability to model complex feature interactions out of the box, TabPFN serves as a state-of-the-art, general-purpose baseline for perturbation-prediction problems.

## 3 Results

### 3.1 Performance Analysis and Optimization of ScAPE Models

To explore the behavior of the ScAPE models under various parameter settings and configurations, we conducted an extensive performance analysis. As in the original challenge, we used the Mean Rowwise Root Mean Squared Error (MRRMSE) computed between true and predicted SLPs for all experiments to evaluate the performance of the models, which is defined as:

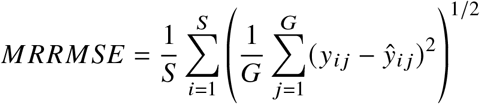

where *S* is the number of samples (rows), *G* is the number of genes (columns), and *y*_*i j*_ and *ŷ*_*i j*_ are the actual and predicted SLP values, respectively, for sample *i* and gene *j* .

First, in order to see if there are groups of genes more informative than others, we evaluated the models using a varying numbers of input genes, selecting the top N most variable genes. Table 1A presents selected configurations and their corresponding performance metrics for the multi-task ScAPE approach, which outperformed the single-task variant.

**Table 1:** Summary of the performance analysis of the multi-task ScAPE model under various parameter configurations and experimental conditions. A) Evaluation across varying numbers of input genes. B) Comparison of ensembling vs. non-ensembling approaches. C) Impact of loss weight configurations: weights of the loss function balancing SLP and LFC error. Best results for each approach and cell type are marked in **bold**. SLPo-w = signed− *log*_10_(p-values) output weight. LFCo-w = *log*_2_ fold changes output weight. Results are evaluated using the Mean Rowwise Root Mean Squared Error (MRRMSE) metric.

| Approach |  | NK | CD4<sup>+</sup> | CD8<sup>+</sup> | T Regulatory | Average |
| --- | --- | --- | --- | --- | --- | --- |
| A<br>Number of Genes | 2 genes | 0.607 | 0.748 | 0.457 | 0.462 | 0.569 |
|  | 8 genes | 0.589 | 0.711 | 0.466 | 0.458 | 0.556 |
|  | 64 genes | **0.577** | **0.689** | **0.445** | **0.450** | **0.540** |
|  | 10,000 genes | 0.591 | 0.700 | 0.456 | 0.454 | 0.550 |
| B<br>Ensemble? | Yes | **0.577** | **0.689** | **0.445** | **0.450** | **0.540** |
|  | No | 0.598 | 0.708 | 0.456 | 0.461 | 0.556 |
| C<br>multi-task Weights | SLPo-w1.0<br>LFCo-w0.0 | **0.575** | 0.699 | **0.444** | **0.449** | 0.542 |
|  | SLPo-w0.8<br>LFCo-w0.2 | 0.577 | **0.689** | 0.445 | 0.450 | **0.540** |
|  | SLPo-w0.5<br>LFCo-w0.5 | 0.583 | 0.697 | 0.452 | 0.456 | 0.547 |
|  | SLPo-w0.2<br>LFCo-w0.8 | 0.592 | 0.722 | 0.451 | 0.457 | 0.555 |
|  | SLPo-w0.0<br>LFCo-w1.0 | 0.637 | 0.807 | 0.464 | 0.478 | 0.596 |

To assess the effect of input dimensionality, we trained models using between 8 and over 10,000 genes. When fewer than 8 genes were provided, accuracy fell sharply, indicating that even modeling the dominant non-response pattern exhibited by most genes requires a minimal feature set. Above this threshold, however, adding three orders of magnitude more genes yielded very little improvement because the training loss remains dominated by genes that do not change under treatment. As a result, the model becomes highly accurate at predicting non-responsive genes but lacks sensitivity for the rare, low p-value differential-expression events that capture true drug effects. The fact that random gene selections perform as well as top-most variable genes underlines the need for better metrics that puts more emphasis on smaller subsets of responsive genes, for example, by up-weighting low-p-value, differentially expressed genes so that the thousands of largely unchanging genes no longer drown out errors on those responsive transcripts.

Additionally, we compared single-fold predictions with an ensemble constructed as the median of the five inner-loop folds (Table 1B). Across every cell type the ensemble lowered the MRRMSE on SLP by 2% to 4%, indicating that pooling the diverse hypotheses learned in each fold yields more stable and accurate predictions.

The multi-task ScAPE network jointly predicts the LFC and SLP for every gene. Its training objective is a weighted sum

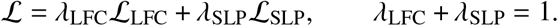

We explored this trade-off by experimenting with different weight configurations, ranging from 1.0 to 0.0 for each output, balancing their contributions to identify the optimal configuration. Our analysis revealed that the model achieved strong performance when both SLP and LFC outputs were considered, with the configuration `SLPo-w0.8 /LFCo-w0.2` yielding the lowest average MRRMSE (Table 1C). This near-balanced regime allows the LFC term to guide the model on the overall scale of expression changes, while the SLP term focuses learning on genes that truly react to each treatment. As a result, the model maintains sensitivity to real biological signals without being overwhelmed by non-responsive genes. On average, the multi-task `SLPo-w0.8 /LFCo-w0.2` achieved lower MRRMSE than the single-task setting, with the largest decrease of 0.01 MRRMSE observed in CD4^+^ T cells.

In summary, ensembling reduces fold-specific variance and improves generalization, and applying a multi-task loss, achieves a good trade-off between learning the general structure of gene expression and focusing on biologically meaningful perturbation effects. This balance translates into improved performance under a metric that averages prediction errors across all genes in each contrast.

### 3.2 Comparative Evaluation of Perturbation Prediction Models

We evaluated the performance of several state-of-the-art architectures, including single-task ScAPE, multi-task ScAPE, NN retraining with pseudolabels, Py-Boost, and TabPFN, using the experimental protocol described in Section 2.3 to ensure comparability. Table 2 summarizes the MRRMSE for each model, alongside the 0-baseline (all values set to zero) and median-baseline (median of the genes grouped by drugs) approaches.

**Table 2:** Performance comparison of various models, including single-task ScAPE, multi-task ScAPE, NN retraining with pseudolabels, Py-Boost, and TabPFN, evaluated using the MRRMSE metric. Baseline approaches (0-baseline and median-baseline) are included for reference. The best results for each cell type are marked in **bold**.

| Approach | NK Cells | CD4<sup>+</sup> T cells | CD8<sup>+</sup> T cells | T Reg. Cells | Average |
| --- | --- | --- | --- | --- | --- |
| single-task ScAPE<sup>2</sup> | **0.573** | 0.697 | 0.445 | 0.451 | 0.542 |
| multi-task ScAPE<sup>2</sup> | 0.577 | **0.689** | 0.445 | 0.450 | **0.540** |
| NN retraining with pseudolabels<sup>3</sup> | 0.657 | 0.824 | 0.511 | 0.518 | 0.628 |
| Py-Boost<sup>4</sup> | 0.596 | 0.747 | **0.441** | **0.449** | 0.558 |
| TabPFN<sup>23</sup> | 0.597 | 0.754 | 0.451 | 0.467 | 0.567 |
| 0-baseline | 0.639 | 0.838 | 0.466 | 0.462 | 0.601 |
| median-baseline | 0.636 | 0.789 | 0.573 | 0.588 | 0.647 |

The single-task and multi-task ScAPE models performed similarly overall, with the multi-task variant achieving slightly better average results (MRRMSE of 0.540 versus 0.542). While differences were minimal across most cell types, multi-task ScAPE demonstrated clearer advantages in CD4^+^ T cells (MRRMSE of 0.689 versus 0.697).

The NN retraining with pseudolabels approach, one of the winning models of the challenge, required substantial integration and adaptation to fit our experimental protocol. While we incorporated validation steps and a robust evaluation mechanism absent in the original implementation, this architecture showed inferior performance, with an average MRRMSE error of 0.628—the highest among all models evaluated. Upon further analysis, we identified a key limitation in our adaptation: due to the fine-tuning of our experimental setup, we were unable to provide the full dataset when generating pseudolabels in the first stage. In contrast, the original implementation used the complete dataset to generate higher-quality pseudolabels, which were then used to improve performance in the second stage. This constraint in our pipeline likely contributed to the lower predictive accuracy observed in our results.

The Py-Boost architecture performed slightly worse (MRRMSE of 0.558) than the ScAPE models but consistently outperformed both TabPFN and the NN retraining with pseudolabels. Nevertheless, Py-Boost remained competitive for specific cell types, achieving strong results in CD8^+^ T cells and T regulatory cells, with MRRMSE errors of 0.441 and 0.449, respectively.

For the TabPFN model (MRRMSE of 0.567), we adapted it to our experimental protocol by applying one-hot encoding and training a separate regressor for each gene and cell type. While TabPFN is specifically designed for tabular data, its performance was notably close to that of Py-Boost in NK cells and CD4^+^ T cells. This demonstrates TabPFN’s remarkable adaptability, achieving competitive results even when applied to a data structure outside its primary goal. However, despite this adaptability, simpler approaches like ScAPE and Py-Boost consistently outperformed TabPFN across all cell types.

Baseline comparisons further emphasize the predictive capabilities of the evaluated models. Across all cell types, the 0-baseline and median-baseline approaches consistently underperformed relative to the ML-based models, except in some cell types, where the NN retraining with pseudolabels approach performed similarly to or worse than baseline approaches. Notably, for NK cells and CD4^+^ T cells, where the best MRRMSE scores were 0.573 and 0.689, respectively, the median-baseline yielded substantially higher errors (0.636 and 0.789), and the 0-baseline performed even worse. These results highlight that the models do not merely default to predicting values close to zero; instead, they are at least able to distinguish which genes are responsive and which are not, in a cell-specific manner.

Across all architectures, the lowest error rates were observed for CD8^+^ T cells, while CD4^+^ T cells consistently yielded the highest MRRMSE values. Multi-task ScAPE achieved the best overall performance, closely followed by single-task ScAPE and Py-Boost. Although the average performance of Py-Boost is below ScAPE, it remains competitive, particularly in CD8^+^ T cells and T regulatory cells. These results reinforce the ability of the model to capture meaningful perturbation responses across diverse cell types and clearly outperform statistical baselines, underscoring their potential for robust and generalizable perturbation prediction. Overall, these results validate the effectiveness of ScAPE.

## 4 ScAPE Tool and API

We also provide an open-source implementation of ScAPE, offering both a command-line interface (CLI) for rapid execution and a flexible Python API for advanced customization.

### 4.1 Command-Line and API Workflows

The ScAPE package includes a simple CLI for model training. A typical command initiates training using parquet files containing SLPs and LFCs. Both interfaces expect input data as Pandas DataFrames with a “MultiIndex” of “(cell_type, sm_name)” where “sm_name” are the names of the drugs. Users can specify hyperparameters and define cross-validation sets by holding out specific cell types or drugs.

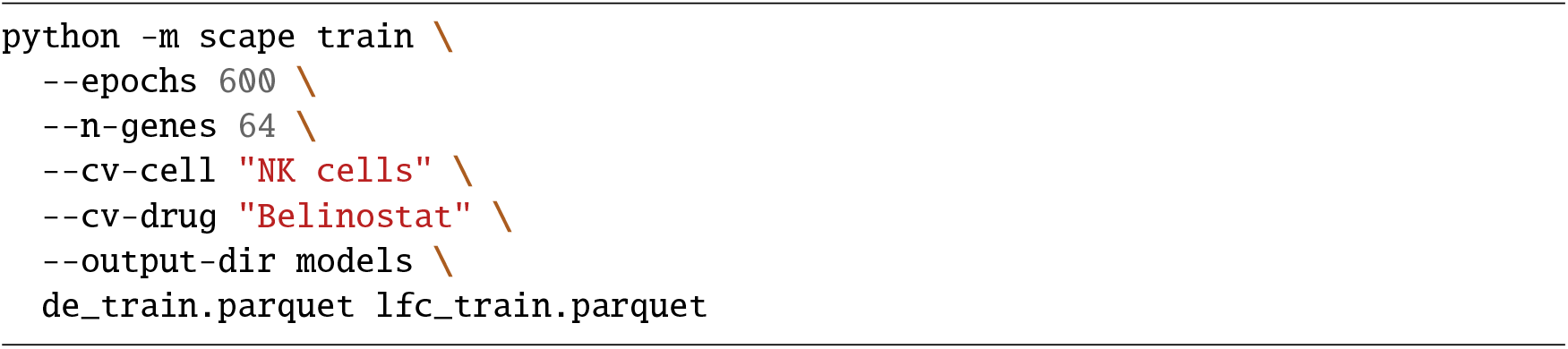

For greater control and integration into larger pipelines, the same workflow can be executed via the Python API, which provides functions for data loading, model instantiation, training, and prediction.

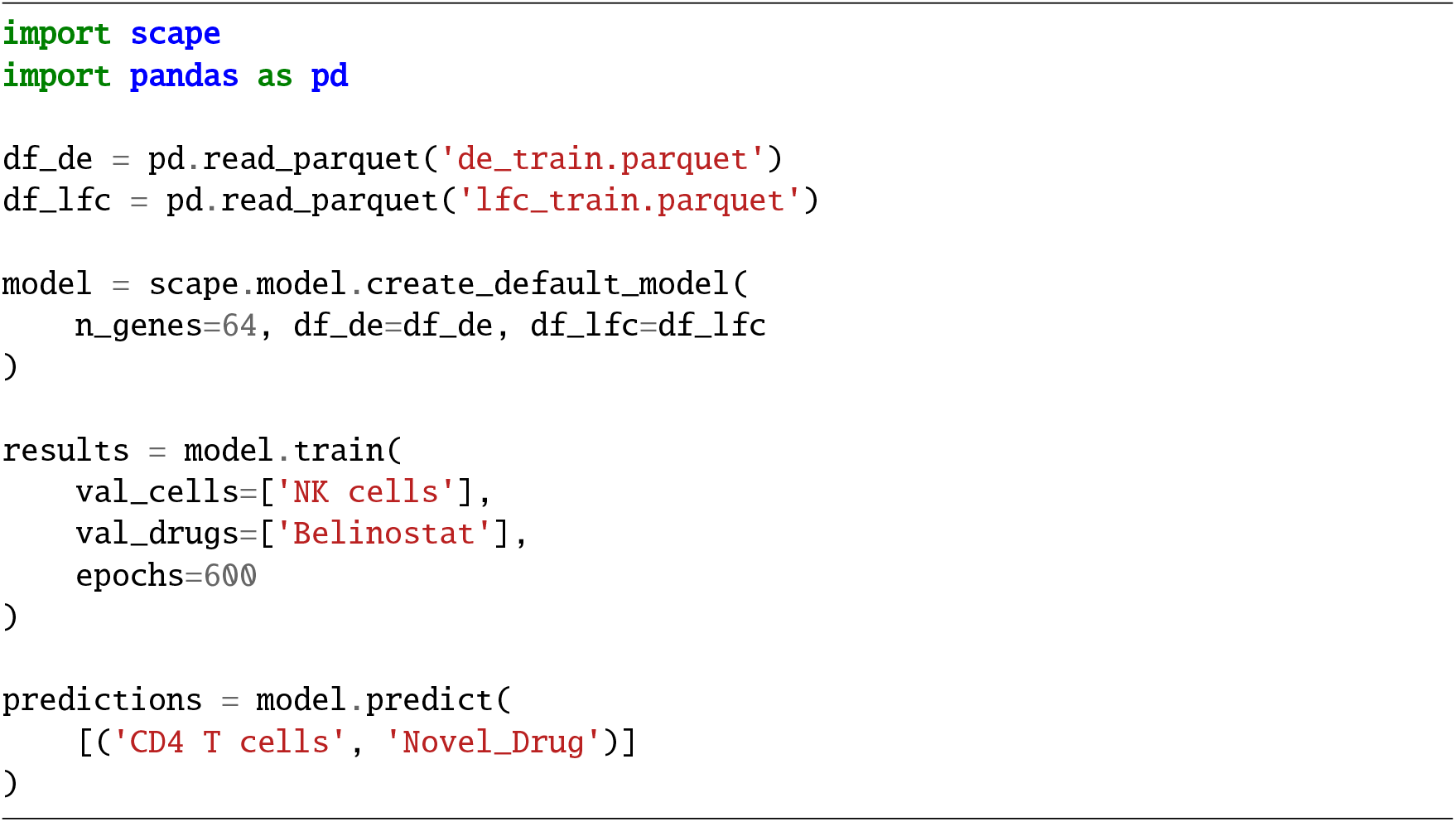

### 4.2 Advanced API Customization

The ScAPE API exposes the underlying model components, allowing for extensive customization of the network architecture, learning objectives, and feature engineering. A key feature is the ability to configure multi-task models that simultaneously predict multiple outputs, such as SLP and LFC. This encourages the network to learn shared representations and can improve generalization.

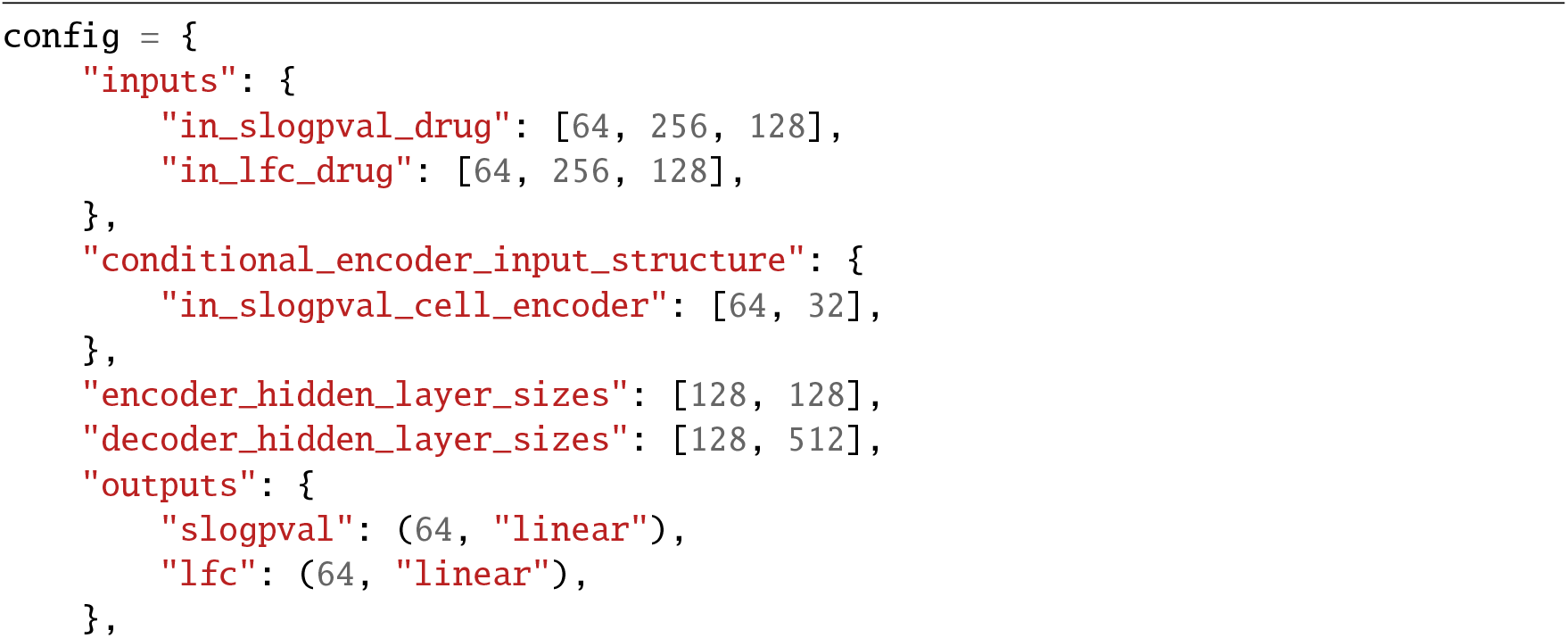
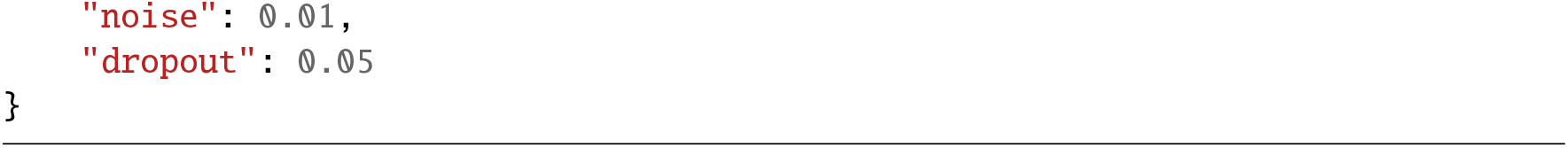

Furthermore, in a multi-task setting, users can prioritize one learning objective over another by assigning different weights to their respective loss functions during model compilation.

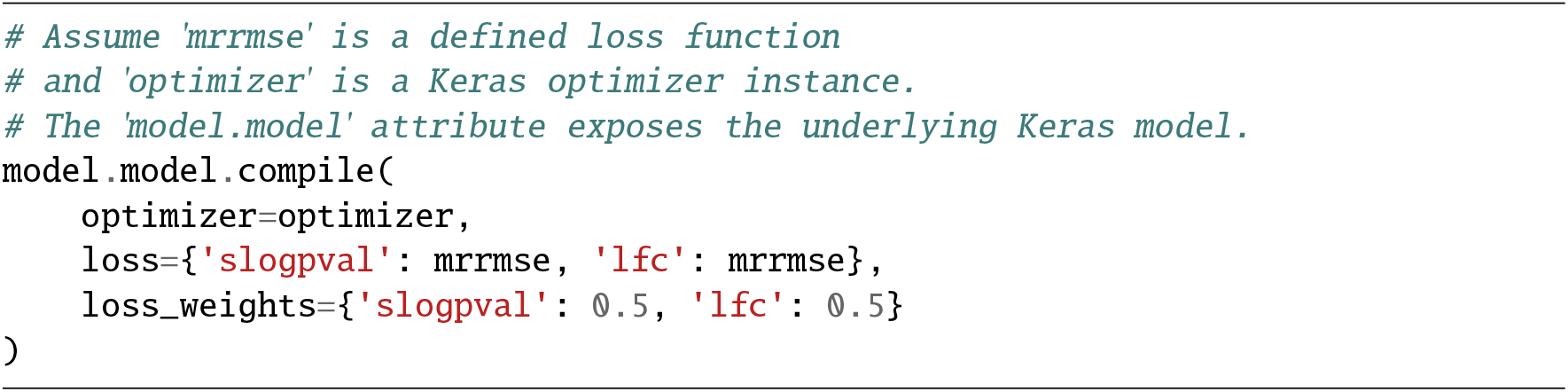

### 4.3 K-Fold Ensemble for Robust Prediction

For enhanced robustness and accuracy, we implement an ensemble of ScAPE models, trained on different subsets of the data. This can be easily achieved by using a K-Fold strategy, generating K trained models on different partitions. This strategy mitigates the risk of overfitting and improves generalization.

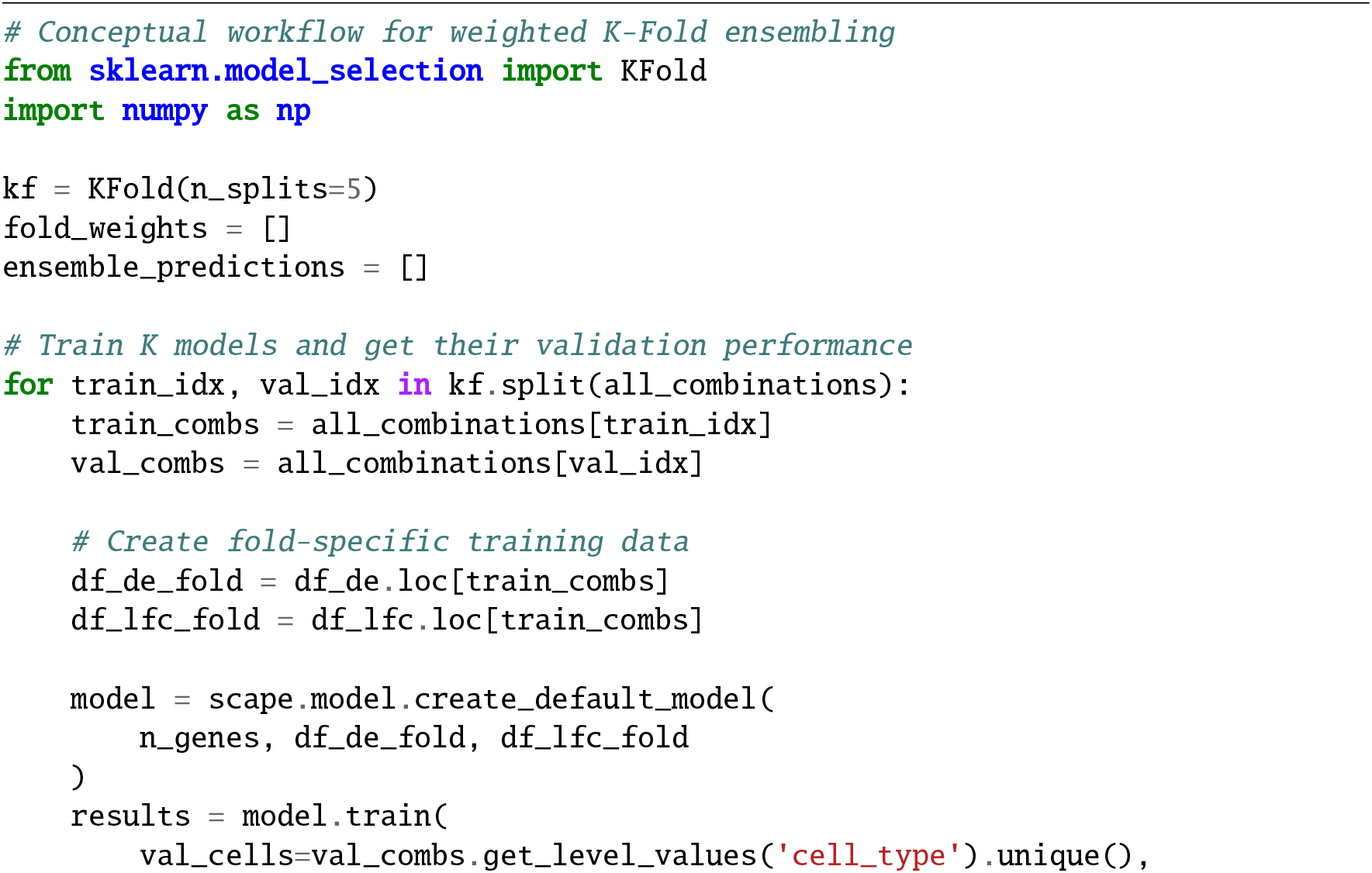
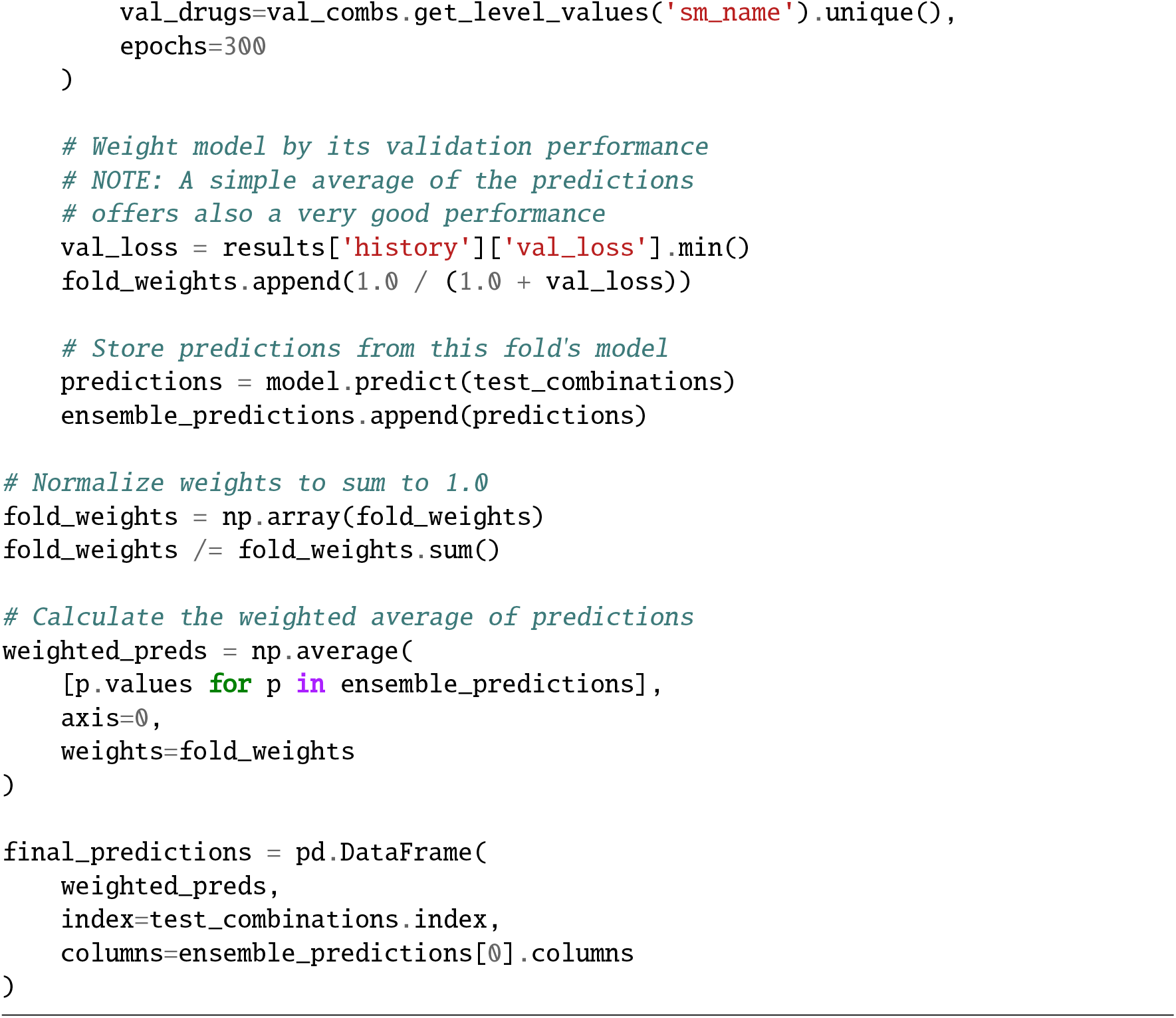

## 5 Discussion

Predicting cellular responses to perturbations remains an important challenge in computational biology, with important implications for drug discovery and personalized medicine. The NeurIPS 2023 perturbation prediction competition provided valuable insights into both the potential and limitations of current approaches.

Building on the competition, our results demonstrate that ScAPE, despite its simplicity, achieves competitive performance. The multi-task variant showed marginal but consistent improvements over the single-task version, which can be attributed to the inherent relationship between p-values and fold changes; by jointly learning both outputs, the model captures their statistical dependencies, with the LFC task serving as an auxiliary regularizer that improves the main target prediction of this challenge, which consists of SLP of the DE test between control and treated cells. A key aspect of ScAPE is the use of median-based features to characterize the drug and cell specific effects on genes. By reducing the high-dimensional gene expression space to just a few features per gene (drug-level and cell-level medians), we transform a complex learning problem into one of modeling biologically meaningful summary statistics. This strategy mirrors successful approaches in collaborative filtering, and simplifies also the strategy of modern methods that also decompose effects into perturbation and context specific covariates^10,11^. The fact that such a simple representation performs competitively suggests that much of the predictive signal can be captured through these population-level summaries.

Our benchmarks using the OP3 dataset show that this approach remains competitive despite its simplicity. While Py-Boost showed strong performance in specific cell types and TabPFN demonstrated remarkable adaptability, despite being a foundation model for tabular data, the overall performance differences between top models were modest. This convergence of performance across diverse architectures suggests we may be approaching fundamental limits imposed by the size of the dataset, the data processing pipeline itself, and the evaluation metrics, even before considering model architecture.

The perturbation challenge used to systematically evaluate the methods relies on upstream computational cell type annotation; errors in identifying cell types, especially for perturbed cells that differ markedly from the unperturbed or basal cell atlases used to train these models^24^, can propagate downstream and degrade training data quality. Furthermore, the final data used for prediction is summarized at the pseudobulk level, averaging away the heterogeneity and subtle population shifts inherent to single-cell resolution. When all models are trained on this lower-resolution signal, it is not surprising to find that the differences between predictive models are minimal. Our enrichment analysis provides some insight into this problem, showing that the genes with the highest prediction error are concentrated in specific biological pathways. These are hallmarks of inflammatory and stress responses, such as TNF-*α* signaling, which often exhibit switch-like activation. The genes within these pathways act as markers for a cellular response, yet the pseudobulking process averages these distinct states into a single, intermediate value that represents neither population accurately. Compounding this biological issue, an unspecific validation metric like MRRMSE on SLP averages statistical significance of differential expressed genes across all 18,211 genes, giving equal weight to thousands of non-responsive genes and the handful of critical response markers. Consequently, the modeling task becomes less about identifying specific biological responses and more about predicting a smoothed average, which helps explain why even sophisticated models struggle to outperform simple baselines under this evaluation scheme.

This observation highlights the importance of designing better metrics for benchmarking perturbation-based models. Beyond the choice of evaluation metric, the objective designed for the OP3 challenge, i.e., the direct prediction of log p-values, is one of the major bottlenecks. An alternative approach could be to predict full single-cell gene expression profiles, where models predict target gene expression given a perturbation and baseline conditions. From such generative predictions, any desired downstream statistical test could be performed post-hoc, providing a statistically grounded and more interpretable output, while also leveraging information from the entire single-cell data without relying on intermediate preprocessing steps such as pseudobulking and computational cell type annotation, as done in the challenge. The combination of these limitations creates a paradoxical situation where models that simply predict the mean expression profile can achieve competitive scores while failing to capture the sparse but biologically critical perturbation responses. This results in an evaluation where architectural complexity offers no clear advantage, as shown in our experiments where models using as few as the top 8 most variable genes performed comparably to those using over 10,000. The training loss remains dominated by non-responsive genes, so the model learns to predict the baseline state accurately but lacks sensitivity for the true biological signals.

In conclusion, our work demonstrates that a simple yet robust model like ScAPE can achieve top-tier performance in a large-scale, unbiased competition where over 1,000 teams proposed and evaluated different methods, suggesting that future progress in perturbation prediction depends as much on better and more large-scale single-cell perturbational assays as well as on the development of better metrics and evaluation frameworks. New benchmarks^25^ should therefore prioritize more biologically relevant objectives, moving from predicting statistical summaries to full single-cell profiles, and must adopt metrics that capture the sparse but critical gene responses that define cellular states^26^. Within such improved frameworks, simple and efficient models like ScAPE can be used as essential baselines, ensuring that the performance gains of novel architectures are genuine and not artifacts of the evaluation setup.

## Acknowledgements

This work was supported through state funds approved by the State Parliament of Baden-Württemberg for the Innovation Campus Health + Life Science Alliance Heidelberg Mannheim. It was also supported by projects: AI4FOOD-CM (Y2020/TCS6654), FACINGLCOVID-CM (PD2022-004-REACT-EU), INTER-ACTION (PID2021-126521OB-I00 MICINN/FEDER), HumanCAIC (TED2021-131787BI00 MICINN), PowerAI+ (SI4/PJI/2024-00062), and Cátedra ENIA UAM-VERIDAS en IA Responsable (NextGenerationEU PRTR TSI-100927-2023-2). We also thank Daniel Dimitrov for valuable feedback.

## Author Contributions

PRM and MGR conceived the original ideas and developed the initial implementation used for the participation in the “Open Problems - Single-Cell Perturbations” competition, part of the NeurIPS 2023 Competition Track. For this study, SRT, PRM, and MGR contributed equally to extending the method, developing the evaluation framework, and writing the manuscript. RT and AM provided supervision to SRT, while JSR supervised the study and provided additional guidance. All authors contributed to review and editing, and approved the final version of the manuscript.

## Footnotes

1 https://www.kaggle.com/competitions/open-problems-single-cell-perturbations

2 https://github.com/scapeML/scape

3 https://github.com/okon2000/single_cell_perturbations

4 https://github.com/Ambros-M/Single-Cell-Perturbations-2023

